# Aire-dependent genes undergo Clp1-mediated 3’UTR shortening associated with higher transcript stability in the thymus

**DOI:** 10.1101/837880

**Authors:** Clotilde Guyon, Nada Jmari, Francine Padonou, Yen-Chin Li, Olga Ucar, Noriyuki Fujikado, Fanny Coulpier, Christophe Blanchet, David E. Root, Matthieu Giraud

## Abstract

The ability of the immune system to avoid autoimmune disease relies on tolerization of thymocytes to self-antigens whose expression and presentation by thymic medullary epithelial cells (mTECs) is controlled predominantly by Aire at the transcriptional level and possibly regulated at other unrecognized levels. Aire-sensitive gene expression is influenced by several molecular factors, some of which belong to the 3’end processing complex, suggesting they might impact transcript stability and levels through an effect on 3’UTR shortening. We discovered that Aire-sensitive genes display a pronounced preference for short-3’UTR transcript isoforms in mTECs, a feature preceding Aire’s expression and correlated with the preferential selection of proximal polyA sites by the 3’end processing complex. Through an RNAi screen and generation of a lentigenic mouse, we found that one factor, Clp1, promotes 3’UTR shortening associated with higher transcript stability and expression of Aire-sensitive genes, revealing a post-transcriptional level of control of Aire-activated expression in mTECs.

## Introduction

Immunological tolerance is a key feature of the immune system that protects against autoimmune disease by preventing immune reactions against self-constituents. Central tolerance is shaped in the thymus and relies on a unique property of a subset of medullary thymic epithelial cells (mTECs). This subset is composed of mTEChi that express high levels of MHC class II molecules and a huge diversity of self-antigens (Danan-Gotthold, Guyon, Giraud, Levanon, & Abramson, 2016; Kyewski & Klein, 2006). Thus, developing T lymphocytes in the thymus are exposed to a broad spectrum of self-antigens displayed by mTEChi. Those lymphocytes that recognize their cognate antigens undergo either negative selection, thereby preventing the escape of potentially harmful autoreactive lymphocytes out of the thymus, or differentiate into thymic regulatory T cells beneficial for limiting autoreactivity (Cowan et al., 2013; Goodnow, Sprent, Fazekas de St Groth, & Vinuesa, 2005; Klein, Kyewski, Allen, & Hogquist, 2014). Self-antigens expressed by mTEChi include a large number of tissue-restricted antigens (TRAs), so-named because they are normally restricted to one or a few peripheral tissues (Derbinski, Schulte, Kyewski, & Klein, 2001; Sansom et al., 2014). A large fraction of these TRAs in mTEChi are induced by a single transcriptional activator that is expressed almost exclusively in these cells - the autoimmune regulator Aire. Mice deficient for the *Aire* gene exhibit impaired TRA expression in mTEChi, whereas TRA expression remains normal in peripheral tissues of these mice. Consistent with inadequate development of central tolerance, *Aire* knockout (KO) mice develop autoantibodies directed at some of these TRAs, resulting in immune infiltrates in multiple tissues (Anderson, 2002). Correspondingly, loss-of-function mutations in the human *AIRE* gene result in a multi-organ autoimmune disorder known as autoimmune polyglandular syndrome type 1 (Nagamine et al., 1997; Peterson, Pitkänen, SILLANPAA, & Krohn, 2004).

How the expression of thousands of Aire-sensitive self-antigen genes is controlled in mTEChi has been a subject of extensive investigation. Significant progress has been made, notably through the identification of a number of molecular factors that further activate the expression of prototypic Aire-sensitive genes in a model employing cell lines that express Aire ectopically by transfection with a constitutive *Aire* expression vector (Abramson, Giraud, Benoist, & Mathis, 2010; Giraud et al., 2014). These studies revealed a role for relaxation of chromatin in front of the elongating RNA polymerase (RNAP) II by the PRKDC-PARP1-SUPT16H complex (Abramson et al., 2010) and for an HNRNPL-associated release of the stalled RNAPII (Giraud et al., 2014). However, the effect of most of the identified factors on the full set of Aire-sensitive genes in mTEChi is unknown. It remains also uncertain whether these factors partake in a molecular mechanism directly orchestrated by Aire or in a basal transcriptional machinery that would control the expression of Aire-sensitive genes even before Aire is expressed in mTEChi. Lack of knowledge of the precise modus operandi of the identified factors potentially leaves major aspects of promiscuous mTEChi gene expression unknown.

Among the identified factors, seven of them, namely CLP1, DDX5, DDX17, PABPC1, PRKDC, SUPT16H and PARP1, have been reported to belong to the large multi-subunit 3’ end processing complex (de Vries et al., 2000; Shi et al., 2009) which controls pre-mRNA cleavage and polyadenylation at polyA sites (pAs) (Colgan & Manley, 1997). Hence, we asked whether any of these identified factors could influence Aire-sensitive gene expression partially or entirely by the way of an effect on pre-mRNA 3’ end maturation. Deep sequencing approaches have revealed that the vast majority of protein-coding genes in mammal genomes (70 - 79%) have multiple pAs mostly located in 3’UTRs (Derti et al., 2012; Hoque et al., 2013). These genes are subject to differential pA usage through alternative cleavage and polyadenylation directed by the 3’ end processing complex and are transcribed as isoforms with longer or shorter 3’UTRs depending on the pA usage (Tian & Manley, 2013). The 3’ end processing complex is composed of a core effector sub-complex comprising CLP1 (de Vries et al., 2000; Mandel, Bai, & Tong, 2008) and a number of accessory proteins that include DDX5, DDX17, PABPC1, PRKDC, SUPT16H and PARP1 (Shi et al., 2009). Although the individual roles of accessory proteins on differential pA usage remain largely unknown, the core protein Clp1 has been reported to favor proximal pA selection in yeast based on depletion experiments (Holbein et al., 2011). Similarly, increasing levels of Clp1 bound to the 3’ end processing complex was also shown to favor proximal pA selection and shorter 3’UTR isoforms (Johnson, Kim, Erickson, & Bentley, 2011). In contrast, a preference for distal pA selection was reported for higher Clp1 levels in a mouse myoblast cell line based on siRNA Clp1 loss-of-function experiments (Li et al., 2015).

Given the observations from prior work that many of the genes, other than Aire itself, that modulate the expression of Aire-induced genes, are members of the 3’ end processing complex, and that one such member of this complex, Clp1, has been reported to affect pA selection, we speculated that 3’UTR length and regulation might be involved in expression of TRAs in mTEChi. We therefore set out to investigate relationships between Aire sensitivity, 3’ end processing, and pA selection in mTEChi.

## Results

### Aire-sensitive genes show a preference for short-3’UTR transcript isoforms in mTEChi and in some peripheral tissues

To assess the proportion of long and short-3’UTR transcript isoforms in mTEChi, we selected the genes that harbor potential proximal alternative pAs in their annotated 3’UTRs according to the PolyA_DB 2 database which reports pAs identified from comparisons of cDNAs and ESTs from a very large panel of peripheral tissues (Lee, Yeh, Park, & Tian, 2007). For each gene, the relative expression of the long 3’UTR isoform versus all isoforms could be defined as the distal 3’UTR (d3’UTR) ratio, i.e., the expression of the region downstream of the proximal pA (d3’UTR) normalized to the upstream region in the last exon (Figure 1A and **Figure 1–source data 1**). To determine whether the Aire-sensitive genes exhibit a biased proportion of long and short-3’UTR isoforms in mTEChi, we first performed RNA deep-sequencing (RNA-seq) of mTEChi sorted from WT and *Aire*-KO mice in order to identify the Aire-sensitive genes, i.e., those upregulated by Aire (Figure 1B). We then compared the distribution of d3’UTR ratios in Aire-sensitive versus Aire-neutral genes. We found a significant shift towards smaller ratios, revealing a preference of Aire-sensitive genes for short-3’UTR isoforms in mTEChi (Figure 1C, Figure 1–figure supplement 1A and **Figure 1– source data 2**). Since a much larger proportion of Aire-sensitive genes than Aire-neutral genes are known to be TRA genes, we asked whether the preference of genes for short-3’UTRs was more aligned with Aire-sensitivity or being a TRA gene. To this end, we compared the d3’UTR ratios between the TRA and non-TRA genes as defined in reference (Sansom et al., 2014) in the subsets of Aire-sensitive and neutral genes. In these mTEChi, the short-3’UTR isoform preference was observed preferentially in Aire-sensitive genes regardless of whether or not they were TRA genes (Figure 1D).

**Figure 1.**
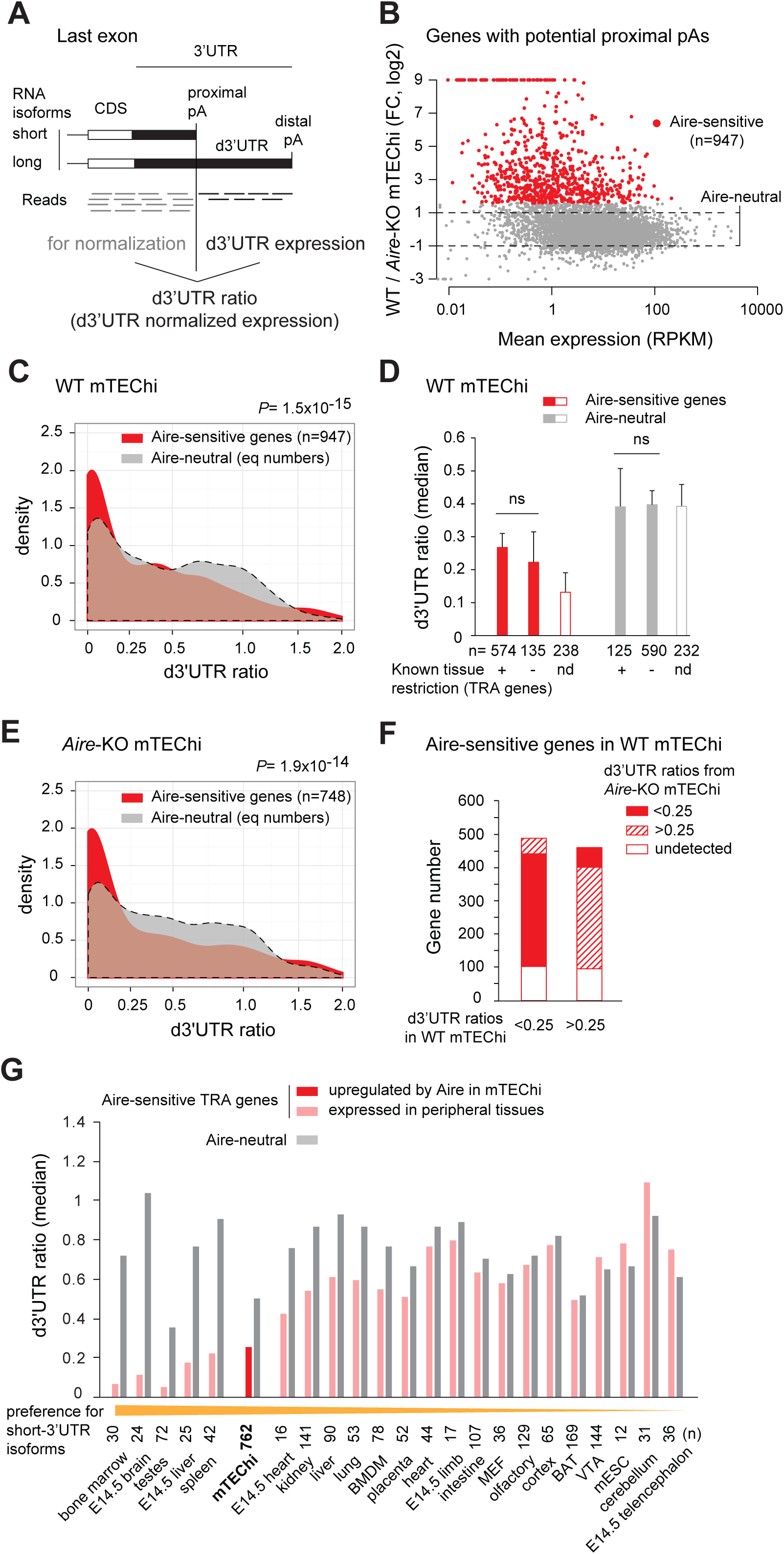
Preference of Aire-sensitive genes for short-3’UTR transcript isoforms in mTEChi and in some peripheral cells. (**A**) Schematic of pA usage and 3’UTR isoform expression. 3’ ends of RNA isoforms of a hypothetical gene are shown. Usage of the proximal pA results in a reduced proportion of the long 3’UTR isoform, estimated by the d3’UTR ratio. (**B**) RNA-seq differential expression (fold-change) between WT and *Aire*-KO mTEChi sorted from a pool of 4 thymi. Red dots show genes upregulated by threefold or more (Z-score criterion of *P*< 0.01) (Aire-sensitive). Genes between the dashed lines have a change in expression less than twofold (Aire-neutral). (**C**) Densities of d3’UTR ratios of Aire-sensitive genes upregulated by Aire in mTEChi and of Aire-neutral genes; equal number (n=947) of neutral genes included, asinh scale. (**D**) Median of d3’UTR ratios of Aire-sensitive and neutral genes depending on whether their peripheral expression is tissue-restricted, or not. Genes whose classification is not established are called “not determined” (nd) and represented by an open box. (**E**) Densities of d3’UTR ratios of Aire-sensitive and neutral genes in *Aire*-KO mTEChi; equal number (n=748) of neutral genes included, asinh scale. (**F**) Proportion of Aire-sensitive genes with d3’UTR ratios <0.25 or >0.25 in *Aire*-KO mTEChi among those with d3’UTR ratios <0.25 or >0.25 in WT mTEChi. (**G**) Median of d3’UTR ratios calculated from RNA-seq data for Aire-sensitive genes with tissue-restricted expression and Aire-neutral genes in mTEChi and 22 mouse tissues. Duplicate reads were discarded to allow more accurate dataset comparison. Cell types were arranged in descending order based on their preference for short-3’UTR isoforms assessed by dividing the median of d3’UTR ratios of Aire-neural genes (gray) by the one of Aire-sensitive genes (pink or red) in each cell type.

To discriminate whether the preference for short-3’UTR isoforms in the Aire-sensitive genes was directly associated with the process of Aire’s induction of gene expression or rather was a feature of Aire-sensitive genes preserved in the absence of Aire, we analyzed the proportion of long 3’UTR isoforms for the Aire-sensitive genes in *Aire*-KO mTEChi. We note that by definition these genes are all expressed at lower levels in the absence of Aire but that most are still expressed at levels sufficient to determine 3’UTR isoform ratios. We observed a preference for the smaller ratios (Figure 1E, Figure 1–figure supplement 1B and **Figure 1– source data 2**) in Aire-sensitive versus neutral genes in the *Aire*-KO cells. We further noted that the majority (∼90%) of Aire-sensitive genes with small d3’UTR ratios (<0.25) in WT mTEChi were also characterized by small d3’UTR ratios in *Aire*-KO mTEChi (Figure 1F and Figure 1–figure supplement 1C), Together, these observations showed that the short-3’UTR isoform preference of Aire-sensitive genes in mTEChi was specific to those genes responsive to Aire, whether or not Aire was actually present.

To determine if genes sensitive to Aire exhibit a preference for short-3’UTR isoforms in their normal tissue of expression, we collected RNA-seq datasets corresponding to a variety of tissues (Shen et al., 2012; van den Berghe et al., 2013; Warren et al., 2013), selected the Aire-sensitive genes characterized by a tissue-restricted expression as identified by the SPM (Specificity Measurement) method (Pan et al., 2013) (Figure 1–figure supplement 1D,E), the Aire-neutral-genes, and compared the proportion of their 3’UTR isoforms in the individual tissues. We found variable preference for short-3’UTR isoforms across peripheral tissues, ranging from none to high, with the mTEChi result falling in the high end of this range (Figure 1G and Figure 1–figure supplement 1F). This finding suggests that factors underlying the preference of Aire-sensitive genes for short-3’UTR isoforms are not necessarily restricted to mTEChi.

### The 3’ end processing complex is preferentially located at proximal pAs of Aire-sensitive genes in AIRE-negative HEK293 cells

We sought to determine whether the preference for short-3’UTR isoforms of Aire-sensitive genes observed in *Aire*-KO mTEChi and conserved upon upregulation by Aire, is associated with a preferred proximal pA usage driven by the 3’ end processing complex. Current techniques dedicated to localize RNA-binding proteins on pre-mRNAs, e.g., ultraviolet crosslinking and immunoprecipitation (CLIP)-seq (König, Zarnack, Luscombe, & Ule, 2012), need several millions of cells, precluding their use with primary *Aire*-KO or WT mTEChi for which only ∼30,000 cells can be isolated per mouse. To circumvent this issue and since we showed (above) that the preference for short-3’UTR isoforms of Aire-sensitive genes was not exclusive to mTEChi, we used the HEK293 cell line. HEK293 cells are (i) negative for AIRE expression, (ii) responsive to the transactivation activity of transfected Aire (Abramson et al., 2010; Giraud et al., 2012), and (iii) have been profiled for the RNA binding of the 3’ end processing components by Martin et al. by PAR-CLIP experiments (Martin, Gruber, Keller, & Zavolan, 2012). Similarly to what we found in WT and *Aire*-KO mTEChi, we observed in *Aire*-transfected and Ctr-transfected HEK293 cells significant lower d3’UTR ratios for genes identified by *Aire* transfection to be Aire-sensitive versus Aire-neutral genes (Figure 2A and Figure 2–figure supplement 1A). It should be noted that many Aire-neutral genes featured moderately lower d3’UTR ratios in *Aire*-transfected HEK293 cells than Ctr-transfected cells, as a possible effect of Aire itself on an overall 3’UTR shortening in these cells, but this did not produce the large proportion of very small d3’UTR ratios (<0.2) that were exhibited by the Aire-sensitive genes in *Aire* or Ctr-transfected HEK293 cells. Within HEK293 cells, localization of the 3’end processing complex at proximal or distal pAs was performed by analyzing the binding pattern of CSTF2, the member of the core 3’ end processing complex that has been reported to exhibit the highest binding affinity for the maturing transcripts at their cleavage sites close to pAs (Martin et al., 2012). We first validated for Aire-neutral genes that lower d3’UTR ratios correlated with the preferential location of the 3’ end processing complex at proximal pAs (Figure 2–figure supplement 1B). Then we compared the location of the complex between the Aire-sensitive and neutral genes, and found a dramatic preference for proximal pAs versus distal pAs at Aire-sensitive genes (Figure 2B), showing that the 3’ end processing complex was already in place at proximal pAs of Aire-sensitive genes before Aire expression was enforced in HEK293 cells.

**Figure 2.**
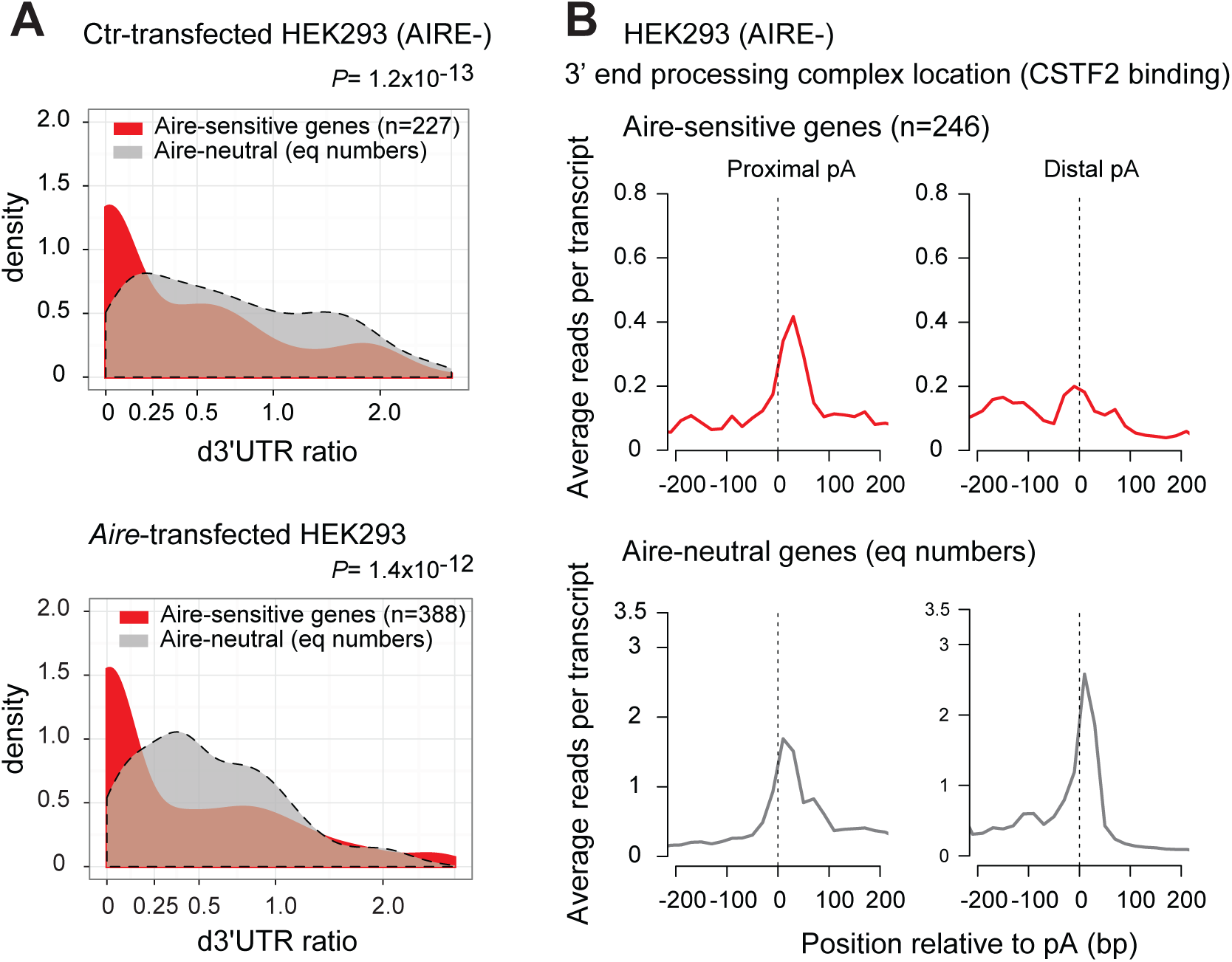
Increased binding of the 3’end processing complex at proximal pAs of Aire-sensitive genes in HEK293 cells. (**A**) Densities in Ctr-transfected HEK293 cells (AIRE-) (*Top*) and *Aire*-transfected HEK293 cells (*Bottom*) of d3’UTR ratios from RNA-seq data of Aire-sensitive and neutral genes identified after *Aire* transfection; equal numbers of neutral genes included (n=227 and n=388, respectively), asinh scale. (**B**) Average density of reads from PAR-CLIP analyses in HEK293 cells (AIRE-) of CSTF2 protein as a marker of the 3’ end processing complex, in the vicinity of proximal and distal pAs of Aire-sensitive and neutral genes. Equal number (n=246) of neutral genes included.

### CLP1 promotes 3’UTR shortening and higher expression at Aire-sensitive genes in HEK293 cells

Since the preference for short-3’UTR isoform expression of Aire-sensitive genes is associated with the increased binding of the 3’ end processing complex to proximal pAs in AIRE-negative HEK293 cells, we asked whether factors that belong to the large 3’ end processing complex (de Vries et al., 2000; Shi et al., 2009) could account for the short-3’UTR transcript isoform preference of Aire-sensitive genes in these cells and affect the expression of these genes. Among the members of the core and accessory subunits of the 3’ end processing complex, the cleavage factor CLP1 (core) and also DDX5, DDX17, PABPC1, PRKDC, SUPT16H and PARP1 (accessory) have been reported to control Aire-sensitive gene expression in an Aire-positive context (Giraud et al., 2014) with the possibility that the effect of some of these factors could result from their action on the basal expression of Aire-sensitive genes and therefore be observed in the absence of Aire. In addition to these candidate factors, we also tested the effect of HNRNPL, which although not part of the 3’ end processing complex, has been shown to regulate 3’ end processing of some human pre-mRNAs (Millevoi & Vagner, 2010) and control Aire transactivation function in mTEChi (Giraud et al., 2014).

First, to determine whether any of the candidate factors could be involved in 3’UTR shortening per se, we carried out short hairpin (sh)RNA-mediated interference in AIRE-negative HEK293 cells and generated expression profiles using Affymetrix HuGene ST1.0 microarrays. These arrays typically include one or two short probes per exon, and for approximatively 35% of all genes with potential proximal pAs they also include at least two short probes in the d3’UTR region. In spite of the limited d3’UTR coverage and lower accuracy of hybridization measurements based on two short probes only, we found these arrays adequate to detect 3’UTR length variation at the genome-scale level. For each gene with a microarray-detectable d3’UTR region, the d3’UTR isoform ratio was calculated by dividing the measured d3’UTR expression by the whole-transcript expression based on all short probes mapping to the transcript. Comparison of the percentages of genes exhibiting a significant increase or decrease of the d3’UTR ratios upon knockdown of a candidate gene versus the control LacZ gene, was used to evaluate the specific impact of the candidate on 3’UTR isoform expression. For each candidate gene, we measured the knockdown efficiency of, typically, 5 different shRNAs and selected the 3 most effective (**Supplementary File 1**). With a threshold of at least 2 shRNAs per gene producing a significant excess of genes with increased d3’UTR ratios, we found that CLP1, DDX5, DDX17, PARP1 and HNRNPL contributed to 3’UTR shortening in HEK293 cells (Figure 3A and Figure 3–figure supplement 1). As a control, we also performed knockdown of CPSF6, a core member of the 3’ end processing complex, that has been consistently reported to promote general 3’UTR lengthening (Li et al., 2015; Martin et al., 2012). As expected, knockdown of *CPSF6* revealed a skewed distribution towards decreased d3’UTR ratios (Figure 3–figure supplement 2).

**Figure 3.**
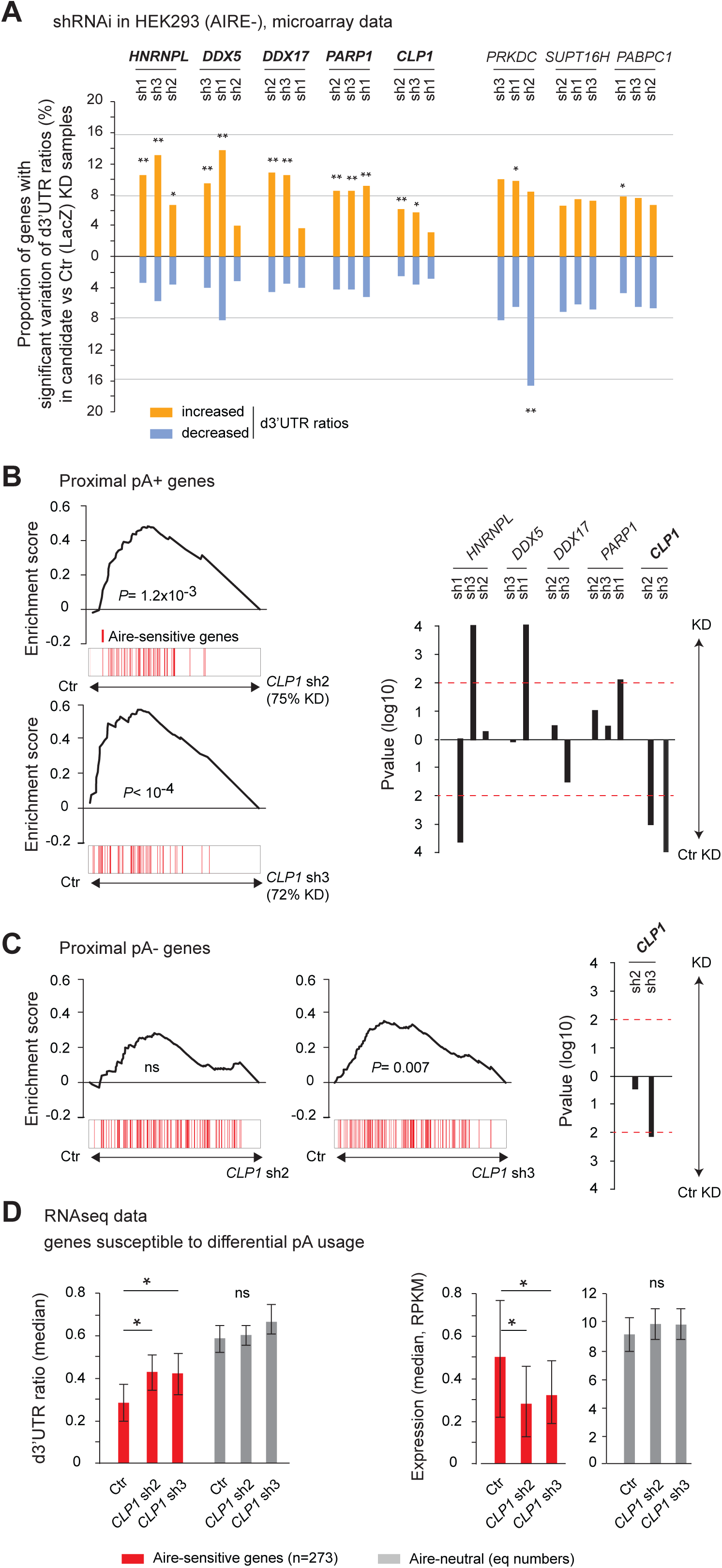
CLP1 controls the expression of Aire-sensitive genes with proximal pAs and their shortening in HEK293 cells. (**A**) Individual probe-level analysis of microarray data from knockdown and control HEK293 cells. Vertical bars represent the proportion of genes with a significant increase or decrease of d3’UTR ratios in the candidate vs. Ctr (LacZ) knockdown samples. * *P*< 10^-4^, ** *P*< 10^-9^ (Chi-squared test). (**B**),(**C**) Gene Set Enrichment Analysis of Aire-sensitive genes among Ctr (LacZ) vs *CLP1* KD ranked expression datasets of all genes with (**B**) or without (**C**) potential proximal pAs in their 3’UTRs as identified in the PolyA_DB 2 database. Significance and direction of the enrichment is shown for each hit shRNA, as well as *P* value thresholds of 0.01 by horizontal red dashed lines (*Right*). (**D**) Median of d3’UTR ratios (*Left*) and expression values (*Right*) from RNA-seq data of Aire-sensitive and neutral genes in HEK293 cells infected by lentiviruses containing *CLP1* hit shRNAs or the Ctr (LacZ) shRNA; equal number (n=273) of neutral genes included, error bars show the 95% confidence interval of the medians. * *P*< 0.05.

Then, we sought among the candidate factors that had an effect on 3’UTR shortening, those that also selectively impacted the basal (in absence of Aire) expression of the Aire-sensitive genes possessing proximal pAs by analyzing microarray whole-transcript expression in control (Ctr) and knockdown HEK293 cells. For each candidate factor, we computed the rank of differential expression of all genes in Ctr versus HEK293 cells knockdown for the tested candidate, and found that *CLP1* was the only gene whose knockdown by two distinct shRNAs led to a significant reduction of the expression of Aire-sensitive genes (Figure 3B), focusing our subsequent study on CLP1. The effect of *CLP1* knockdown on Aire-sensitive genes lacking annotated proximal pAs was much less than for the genes with proximal pAs, but a smaller effect on the annotated single-pA genes was observed for one of two CLP1 shRNAs (Figure 3C). This might simply be because some genes tagged as “proximal pA-“ in the incomplete PolyA_DB 2 database possessed undiscovered proximal pAs in mTEChi and were therefore responsive to the action of CLP1. Our findings regarding the differential response of pA+ and pA-genes to CLP1 loss-of-function suggested that the effect of CLP1 on Aire-sensitive gene expression in HEK293 cells was dependent on the potential of Aire-sensitive genes to switch from using a distal to a proximal pA site.

To assess the effect of CLP1 on 3’UTR shortening at Aire-sensitive genes, we performed RNA-seq experiments in Ctr and *CLP1* knockdown HEK293 cells. Comparison of the 3’ end profiles revealed a statistically significant increase of the median d3’UTR ratios of Aire-sensitive genes for both *CLP1* hit shRNAs (Figure 3D, *Left*), showing that CLP1 is able to promote 3’UTR shortening at Aire-sensitive genes in HEK293 cells. The d3’UTR ratios of Aire-sensitive genes after *CLP1* knockdown did not reach those of Aire-neutral genes which could simply be due to remaining *CLP1* in these cells following knockdown (measured knockdown as 75% and 72% for shRNA 2 and 3, respectively) or could indicate that additional non-CLP1-dependent factors also contribute to the difference in d3’UTR ratios of Aire-sensitive versus Aire-neutral genes. In contrast to Aire-sensitive genes, no significant shift in the d3’UTR ratios of Aire-neutral genes could be detected in Ctr versus *CLP1* knockdown HEK293 cells, consistent with the microarray results (Figure 3A) indicating that only a small proportion of all genes in the genome with microarray-detectable d3’UTR regions underwent 3’UTR length variation after *CLP1* knockdown. Finally, using RNA-seq data, we confirmed the effect of *CLP1* knockdown on the selective reduction of the expression of Aire-sensitive genes that were annotated to have alternative proximal pAs in HEK293 cells (Figure 3D, *Right*).

Together these findings showed that CLP1 is able to promote 3’UTR shortening and increase expression of Aire-sensitive genes with proximal pAs, supporting a linked mechanism between CLP1-promoted 3’UTR shortening and gene expression enhancement at Aire-sensitive genes.

### Clp1 promotes 3’UTR shortening and higher levels of Aire-upregulated transcripts in mTEChi

To test for the *in vivo* impact of Clp1 on 3’UTR length variation of the transcripts upregulated by Aire in mTEChi, we generated lentigenic *Clp1* knockdown mice. Three shRNAs targeting the murine *Clp1* with the highest knockdown efficiency (**Supplementary File 1**) were cloned as a multi-miR construct into a lentiviral vector, downstream of a doxycycline inducible promoter and upstream of the GFP as a marker of activity. Purified concentrated lentiviruses containing this construct and the lentiviral vector expressing the TetOn3G protein were used to microinfect fertilized oocytes, which were reimplanted into pseudopregnant females (Figure 4A). Of the 19 pups that we obtained with integration of both plasmids, two pups were expressing, after doxycycline treatment, the multi-miR in mTEChi and one pup was exhibiting a 60% reduction of *Clp1* mRNA levels in GFP+ (*Clp1* knockdown) versus GFP-(Ctr) mTEChi taken from the same mouse. These two cell populations, *Clp1* knockdown and no-knockdown Ctr mTEChi, were separated by GFP signal by FACS and processed for RNA-seq. We found higher d3’UTR ratios of Aire-sensitive genes in the *Clp1* knockdown mTEChi versus the Ctr mTEChi, whereas no difference was detected for Aire-neutral genes, therefore showing a specific effect of Clp1 on 3’UTR shortening at Aire-sensitive genes in mTEChi (Figure 4B, *Left* and **Figure 4–source data 1**). As in the HEK293 *in vitro* experiments, we observed that the d3’UTR ratios of the Aire-sensitive genes didn’t reach the ratios of the Aire-neutral genes after *Clp1* knockdown, again due either to the incomplete 60% *Clp1* knockdown or to the existence of other non-Clp1-dependent differences.

**Figure 4.**
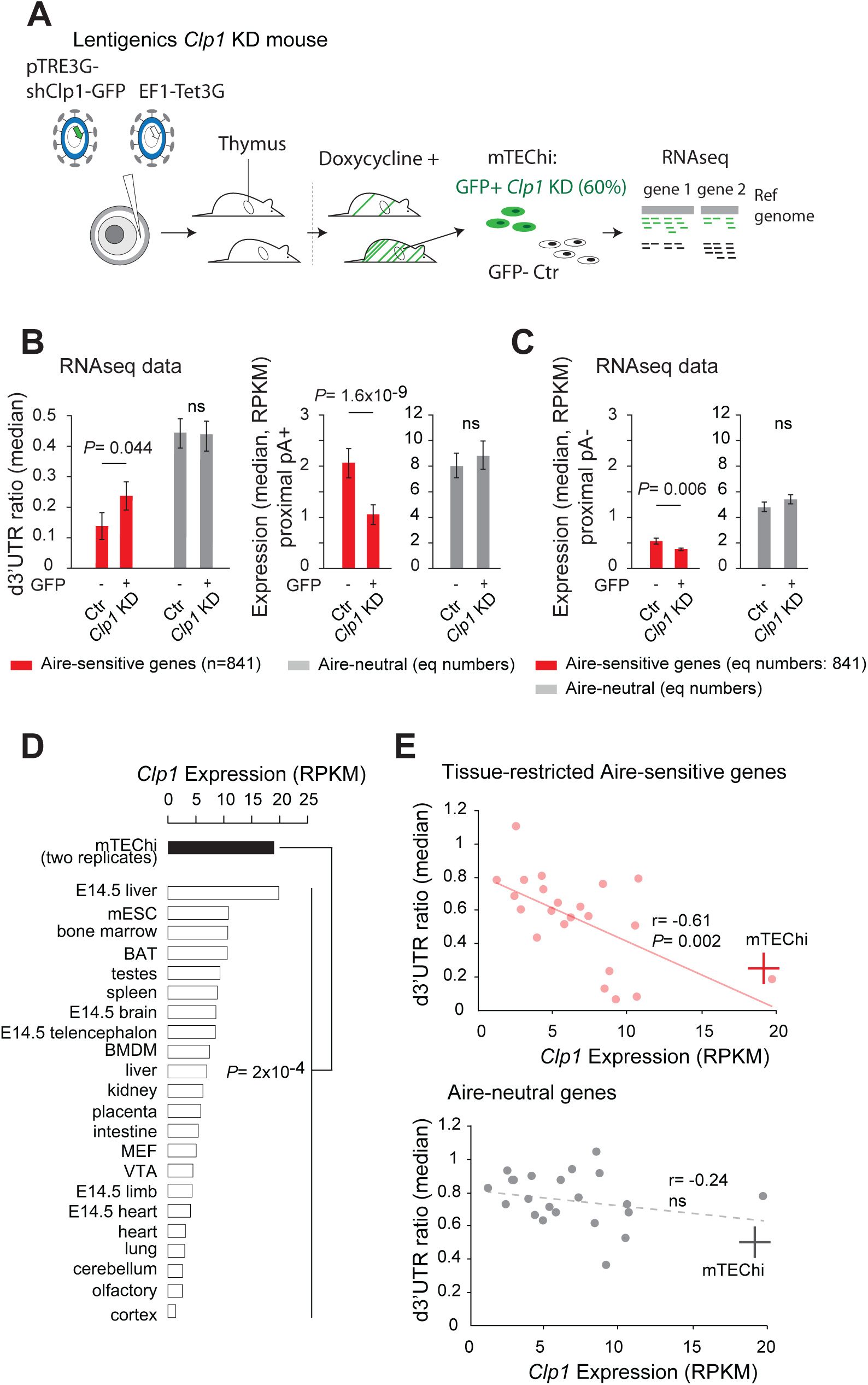
Clp1 controls the expression of Aire-upregulated genes with proximal pAs and their shortening in mTEChi. (**A**) Schematic of the lentigenic knockdown strategy. shRNAs against *Clp1* were transferred to a doxycycline-inducible expression system for microinfection of fertilized oocytes under the zona pellucid. The resulting pups were screened for integration of the constructs and, after treatment by doxycycline, for GFP expression in mTEChi. GFP+ and GFP-mTEChi were sorted and their transcripts profiled by RNA-seq. (**B**) Median of d3’UTR ratios (*Left*) and expression values (*Right*) from RNA-seq data of Aire-sensitive and neutral genes with proximal pAs in GFP+ and GFP-mTEChi from a lentigenic mouse with GFP as a marker of *Clp1* knockdown activity; equal numbers (n=841) of neutral genes included. (**C**) Expression values of Aire-sensitive and neutral genes without proximal pAs in GFP+ and GFP-mTEChi; numbers of included genes (n=841). (**D**) *Clp1* expression from RNA-seq data of two replicate mTEChi and of 22 mouse tissues; median, log10 scale. BAT stands for brown adipocytes tissue, BMDM for bone marrow derived macrophage, MEF for mouse embryonic fibroblast, mESC for mouse embryonic stem cells and VTA for ventral tegmental area. Duplicate reads were discarded for datasets comparison. (**E**) Median of d3’UTR ratios of Aire-sensitive genes whose expression in the periphery is tissue-restricted and of Aire-neutral genes, relative to the levels of *Clp1* expression (log10 scale) in 22 mouse tissues. mTEChi are represented by a red and gray cross for the Aire-sensitive (*Top*) and neutral genes (*Bottom*), respectively; peripheral cells are represented by pink and gray circles. Significance is reached for the Aire-sensitive genes, *P*= 0.002, Pearson correlation.

We then compared the levels of expression of the Aire-sensitive genes with potential proximal pAs in Ctr versus *Clp1* knockdown mTEChi and found, in contrast to Aire-neutral genes, a significant reduction following *Clp1* knockdown (Figure 4B, *Right*). As observed in HEK293 cells, the effect of *Clp1* knockdown was dampened for the Aire-sensitive genes lacking potential proximal pAs (Figure 4C). We also found that the expression of the genes without proximal pAs was globally reduced in comparison to the genes with proximal pAs, supporting a linked mechanism between proximal pA usage and increased gene expression in mTEChi. Altogether these results showed that Clp1 promotes 3’UTR shortening of the transcripts upregulated by Aire in mTEChi and strongly suggested that it enhances their level of expression through proximal pA usage.

A role for Clp1 in mTEChi was further supported by its higher expression in mTEChi in comparison to a wide variety of tissues for which we collected RNA-seq datasets (Shen et al., 2012; van den Berghe et al., 2013; Warren et al., 2013) (Figure 4D). As a comparison, none of the other candidate factors, Hnrnpl, Ddx5, Ddx17 and Paprp1 that contributed to 3’UTR shortening in HEK293 cells, were over-represented in mTEChi (Figure 4–figure supplement 1). Importantly, we also validated higher expression of Clp1 at the protein level in mTEChi versus their precursor cells, mTEClo (Gäbler, Arnold, & Kyewski, 2007; Hamazaki et al., 2007), cells from the whole thymus, and also the predominant thymus CD45+ leukocyte fraction (Figure 4–figure supplement 2). In addition, a role for Clp1 independent of Aire’s action on genes expression was supported by the observed similar levels of *Clp1* expression in WT and *Aire*-KO mTEChi (Figure 4–figure supplement 3A) and the lack of evidence for Aire and CLP1 interaction (Figure 4–figure supplement 3B). Finally, we found a significant correlation between higher *Clp1* expression and lower d3’UTR ratios across peripheral tissues for the Aire-sensitive TRA genes (Figure 4E, *Top*). In contrast, no such significant correlation was observed for Aire-neutral genes (Figure 4E, *Bottom*). These findings indicate that the effect of Clp1 on 3’UTR shortening at Aire-sensitive genes is independent of Aire’s action on gene expression, general across cell types, and conserved upon upregulation of gene expression by Aire in mTEChi.

### Clp1-driven 3’UTR shortening of Aire-upregulated transcripts is associated with higher stability in mTEChi

To determine whether the effect of Clp1 on 3’UTR shortening of Aire-upregulated transcripts was associated with higher levels of these transcripts in mTEChi through increased stability, we assessed the stability of all transcripts in these cells. We used Actinomycin D (ActD) to inhibit new transcription (Sobell, 1985) and assessed the differences in transcript profiles between treated and untreated mTEChi. We harvested the cells at several timepoints for expression profiling by RNA-seq. Each RNA-seq dataset was normalized to total read counts and we calculated the expression fold-change of each gene in treated versus control samples, therefore reflecting differences in transcript levels in the absence of ongoing transcription. We selected among the Aire-sensitive genes two subsets: (i) those that underwent a higher than 2-fold d3’UTR ratio decrease in Ctr versus *Clp1* knockdown lentigenic mTEChi and, (ii) genes with steady d3’UTR ratios (**Figure 4–source data 1**). Comparison of the two gene sets in ActD-treated versus control samples revealed that the Aire-sensitive genes subject to Clp1-driven 3’UTR shortening were initially comparable in their changes in transcript levels upon transcription inhibition to the changes for Aire-sensitive genes that showed no 3’UTR shortening but then showed increasing preservation of transcript levels at 3h and 6 or 12h, indicating a stabilization of this subset of 3’UTR-shortened transcripts (Figure 5; p=2.1×10^-4^, 1.3×10^-12^, and 5.6×10^-10^ at 3, 6 and 12h, respectively). Therefore, the Clp1-promoted 3’UTR shortening at Aire-sensitive genes in mTEChi is indeed associated with higher transcript stability.

**Figure 5.**
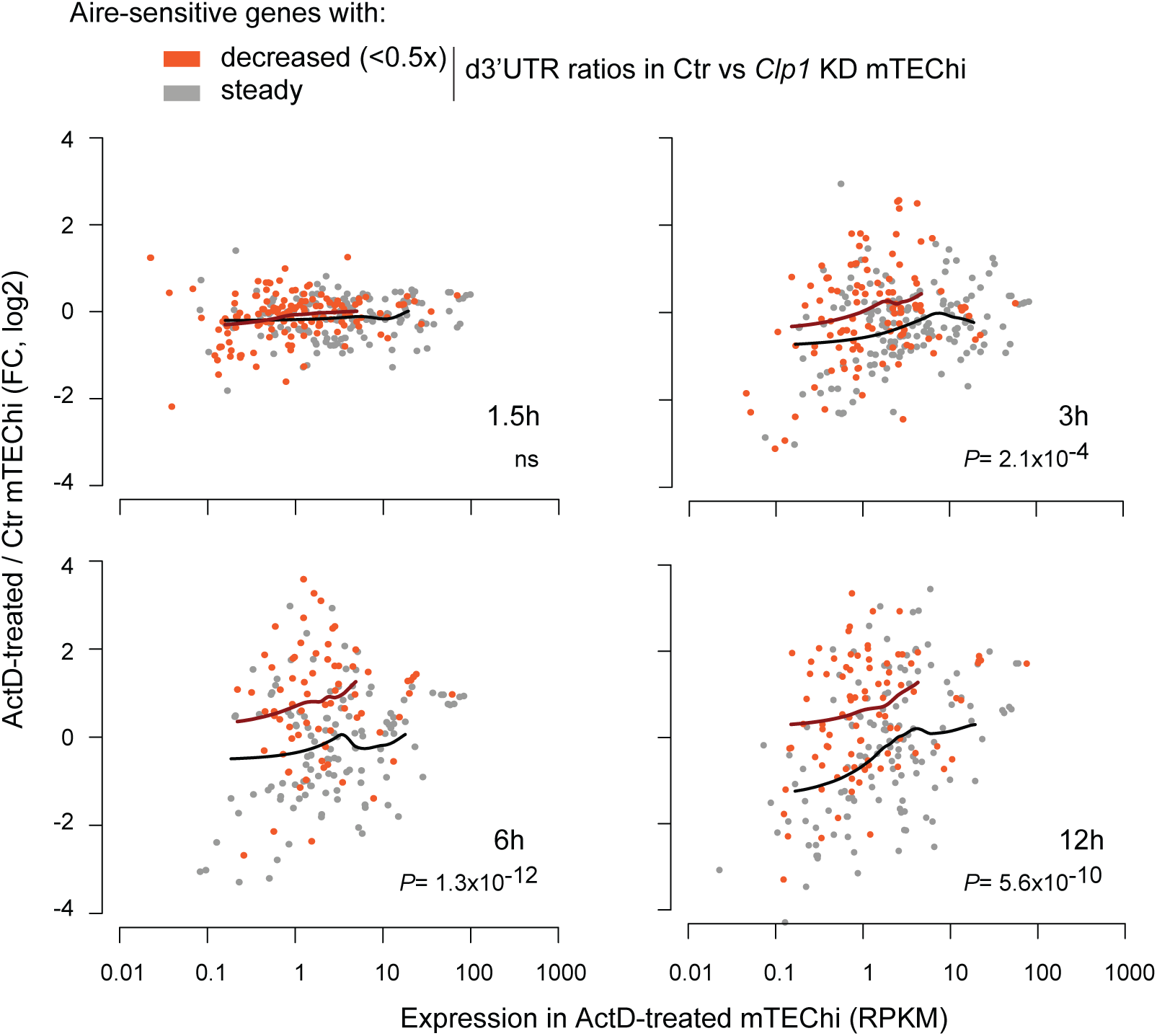
Clp1-driven 3’UTR shortening of the Aire-upregulated transcripts show higher stability in mTEChi. Relative expression of Aire-sensitive genes in ActD-treated (for indicated time durations) vs. control mTEChi depending on whether they undergo Clp1-mediated 3’UTR shortening as identified in *Clp1* lentigenics. Loess fitted curves are shown in dark orange and black for the Aire-sensitive genes with decreased (<0.5x) and steady d3’UTR ratios, respectively. P values are for comparison of expression ratios.

## Discussion

Aire-sensitive genes upregulated by Aire in mTEChi have been shown to be controlled also by a number of Aire’s allies or partners (Abramson et al., 2010; Giraud et al., 2014). Some of these factors have been reported to be part of the large 3’ end processing complex (de Vries et al., 2000; Shi et al., 2009), notably Clp1 which belongs to the core 3’ end processing sub-complex (de Vries et al., 2000; Mandel et al., 2008) and favors early cleavage and polyadenylation in some systems (Holbein et al., 2011; Johnson et al., 2011). In our present study, we demonstrated that Clp1 promotes 3’UTR shortening of the transcripts upregulated by Aire in mTEChi, and that these transcripts are associated with an enhanced stability, revealing a post-transcriptional mechanism whose escape leads to higher levels of expression of Aire-sensitive genes in mTEChi.

Comparison of RNA-seq expression profiles between *Clp1* knockdown and Ctr mTEChi isolated from *Clp1* lentigenic mice showed that Clp1 was able to promote 3’UTR shortening at Aire-sensitive genes specifically, thereby contributing to the preference of short-3’UTR transcript isoforms of Aire-sensitive genes versus Aire-neutral genes in WT mTEChi. We found that this 3’UTR shortening was driven by Clp1 in mTEChi but also in Aire-non-expressing HEK293 cells, in which the level of Aire-sensitive gene expression is weak but still detectable by RNA-seq, thus showing that the effect of Clp1 on Aire-sensitive genes was not restricted to mTEChi nor dependent on Aire’s action, but nonetheless specific to Aire-sensitive genes. Although Clp1 is a main contributor to this described process, additional factors might also be involved.

Clp1 is a ubiquitous protein showing higher expression in mTEChi than in mTEClo, CD45^+^ thymic cells or thymic cells taken as a whole, but also showing higher gene expression in comparison with a large range of peripheral tissue cells. Interestingly, we found that the level of *Clp1* expression was also significantly high in 14.5 embryonic liver cells, cells that have been reported to undergo sustained cellular activation leading to proliferation and maturation into hepatocytes (Kung, Currie, Forbes, & Ross, 2010). Similarities with the pattern of mTEC development (Gray, Abramson, Benoist, & Mathis, 2007; Irla et al., 2008) might point out cell activation as a potent inducer of Clp1 expression. Notably, we showed that higher levels of *Clp1* expression were correlated with higher proportions of short-3’UTR transcript isoforms of tissue-restricted Aire-sensitive genes across peripheral tissues, suggesting the existence of a Clp1-promoted 3’UTR shortening mechanism occurring in a variety of cell types and, among those cell types we surveyed, most pronounced in mTEChi.

The observation that the levels of *Clp1* expression and the distributions of the short-3’UTR transcript isoforms of Aire-sensitive genes were similar between *Aire*-KO and WT mTEChi, strongly suggest that the Clp1-driven 3’UTR shortening mechanism is already in place in mTEChi before Aire is activated. The independence of Aire’s action on gene expression and the effect of Clp1 on 3’UTR shortening is also consistent with the lack of direct interaction found between Aire and Clp1. Although these effects of Aire and Clp1 appear decoupled, it is also apparent as noted that sensitivity to these Aire and Clp1 effects have a high-rate of co-occurrence in the same set of genes. Although we could detect and characterize the effect of Clp1 on the Aire-sensitive genes in mTEChi using annotated pAs from the PolyA_DB 2 database which mainly contains pAs identified from comparisons across peripheral tissues, the precise proportion of Aire-sensitive genes that are subject to differential pA usage and Clp1-driven shortening in mTEChi remains to be precisely defined. Current methods to capture mTEChi-specific pAs using next generation sequencing methods require very large numbers of cells relative to the number of mTEChi (∼30,000) that can be isolated per mouse but emerging methods for accurate pA identification with lower input requirements could make this feasible from such low material quantity (W. Chen et al., 2017).

Clp1 is a member of the core 3’ end processing complex that we showed to be preferentially located at proximal pAs of Aire-sensitive genes in HEK293 cells, indicating that the effect of Clp1 on 3’UTR shortening could result from enhanced recruitment of the 3’ end processing complex to proximal pAs. Similar modes of action, involving members of the core 3’ end processing complex and resulting in transcripts with either shorter or longer 3’UTRs have been described for Clp1 in yeast (Johnson et al., 2011) and Cpsf6 in humans (Gruber, Martin, Keller, & Zavolan, 2012; Martin et al., 2012), respectively. However, the basis for the specificity of Clp1’s effect for the Aire-sensitive genes remains unknown, but note that it does appear to be conserved across cell types. One possibility is that the regulatory elements and associated basal transcriptional machinery at Aire-sensitive genes share conserved features that favor recruitment of Clp1.

3’UTRs have been described as potent sensors of the post-transcriptional repression mediated by miRNAs, resulting in mRNA destabilization and degradation (Bartel, 2009; Jonas & Izaurralde, 2015). In addition to miRNAs, RNA-binding proteins (RBPs) also contribute the regulation of transcript stability depending on the type of cis regulatory elements that they recognize on 3’UTRs, triggering either repression or activation signals (Spies, Burge, & Bartel, 2013). In mTEChi, we found increased stability of the Aire-sensitive transcripts that were subject to Clp1-promoted 3’UTR shortening. Thus, our findings supported an escape from the post-transcriptional repression of short-3’UTR transcripts in mTEChi, leading to enhanced stability and accumulation of these transcripts. Similar observations, resulting in increased transcript levels and higher protein translation, have notably been documented for genes subject to 3’UTR shortening whose long-3’UTR transcript isoforms were targeted by particular miRNAs or classes of RBPs (C. Y. Chen & Shyu, 1995; Guo, Ingolia, Weissman, & Bartel, 2010; Masamha et al., 2014; Mayr & Bartel, 2009; Vlasova et al., 2008).

In addition to impacting transcript stability through the escape of the post-transcriptional repression, short 3’UTRs have also been shown to shift the surface localization of membrane proteins and the functional cell compartment of other types of proteins, in favor of the endoplasmic reticulum (ER) (Berkovits & Mayr, 2015). Thus, one interesting hypothesis for future study is that Clp1-driven 3’UTR shortening at Aire-sensitive genes in mTEChi might not only impact the expression of Aire-dependent self-antigens but also favor their routing to the ER, from where they will be processed and presented, potentially enhancing their presentation to self-reactive T lymphocytes and, subsequently, central tolerance and protection against autoimmune manifestations.

## Materials and Methods

### Mice

*Aire*-deficient mice on the C57BL/6 (B6) genetic background (Anderson, 2002) were kindly provided by D. Mathis and C. Benoist (Harvard Medical School, Boston, MA), and wild-type B6 mice were purchased from Charles River Laboratories. Mice were housed, bred and manipulated in specific-pathogen-free conditions at Cochin Institute according to the guidelines of the French Veterinary Department and under procedures approved by the Paris-Descartes Ethical Committee for Animal Experimentation (decision CEEA34.MG.021.11 or APAFIS #3683 N° 2015062411489297 for lentigenic mouse generation).

### Isolation and analysis of medullary epithelial cells

Thymi of 4-wk-old mice were dissected, trimmed of fat and connective tissue, chopped into small pieces and agitated to release thymocytes. Digestion with collagenase D (1 mg/mL final) (Roche) and DNase I (1 mg/mL final) (Sigma) was performed for 30 min at 37 °C. The remaining fragments were then treated with a collagenase/dispase mixture (2 mg/mL final) (Roche) and DNase I (2 mg/mL final) at 37 °C until a single-cell suspension was obtained. Cells were passed through 70-μm mesh and resuspended in staining buffer (PBS containing 1% FBS and 5 mM EDTA). For isolation of medullary epithelial cells from pooled thymi, an additional step of thymocyte depletion was performed using magnetic CD45 MicroBeads (Miltenyi Biotec). The resuspended cells were incubated for 20 min at 4 °C with the fluorophore-labeled antibodies CD45-PerCPCy5.5 (1:50) (Biolegend), Ly51-PE (1:800) (Biolegend), and I-A/E-APC (1:1,200) (eBioscience). Sorting of mTEChi/lo (CD45^−^PE^−^I-A/E^high/low^) or, for lentigenic mice, of mTEChi +/- for GFP expression, was performed on a FACSAria III instrument (BD Bioscience). For Clp1 staining, cells labeled for membrane antigens were fixed in (3.7%) formaldehyde for 15 min, permeabilized in (0.5%) saponin for 15 min, and incubated with an antibody to Clp1 (1:100) (clone: EPR7181, GeneTex, GTX63930) and an Alexa Fluor 488-conjugated goat polyclonal antibody to rabbit (1:200) (TermoFisher, A11008). Cells were analyzed on an Accuri C6 instrument (BD Bioscience). All compensations were performed on single-color labeling of stromal cells and data analysis was done using the BD Accuri C6 Analysis software.

### Actinomycin D treatment

mTEChi were isolated and sorted (∼4 x 10^5^) from pooled thymi of B6 mice as described above, then treated with actinomycin D (1 µM) in MEM medium for 3, 6 and 12 hours at 37°C. The cells were then harvested and total RNA was isolated by TRIzol extraction (ThermoFisher).

### *Aire*-transient transfections

HEK293 cells were maintained in DMEM high glucose medium complemented with 10% FBS, L-glutamate, sodium pyruvate 1mM, non-essential amino acids and pen/strep antibiotics. The cells were seeded at a density of 600,000 per well in 6-well plates or at 3.5*10^6 per 10-cm2 culture dish. The next day, and depending on the dish format, HEK293 cells were transfected with either 2.5 or 8 ug of the pCMV-Aire-Flag plasmid (or control plasmid) using 7.5 or 32 ul of the TransIT-293 transfection reagent (Mirus). After 48h, cells cultured in 6-well plates were subjected to total RNA extraction for RNA-seq experiments, whereas those in the culture dish were subjected to protein extraction for coimmunoprecipitation.

### Coimmunoprecipitation

Extraction of nuclear proteins and coimmunopreciptation were performed using the Universal Magnetic Co-IP Kit (Active Motif). Briefly, *Aire*-transfected HEK293 cells were lysed with hypotonic buffer and incubated on ice for 15 min. Cell lysates were centrifuged for 30 sec at 14,000 x g, then the nuclei pellets were digested with an enzymatic shearing cocktail for 10 min at 37°C. After centrifugation of the nuclear lysates, the supernatants containing the nuclear proteins were first incubated with specific antibodies for 4 hr, then with Protein-G magnetic beads for 1 hr at 4°C with rotation. After 4 washes, bound proteins were eluted in laemmli/DTT buffer, separated by SDS/PAGE for 40 min at 200 V, transferred to PVDF membranes using the TurboTransfer System for 7 min at 25 V (BioRad) and blocked for 1 hr with TBS, 0.05% Tween, 3% milk. The Western blot detection was done after incubation with primary (2 hr) and secondary antibodies (1 hr). Detection was performed by enhanced chemiluminiscence (ECL). Antibodies used for immunoprecipitation or revelation were: CLP1 (sc-243005, Santa Cruz), Flag-tag M2 (F1804, Sigma), goat IgG control (sc-2028, Santa Cruz), mouse IgG1 control (ab18443, Abcam), and horseradish peroxidase-conjugated anti-mouse IgG light chain specific (115-035-174, Jackson ImmunoResearch).

### shRNA-mediated knockdown

Specific knockdown of *CLP1*, *HNRNPL*, *DDX5*, *DDX17*, *PARP1*, *PRKDC, SUPT16H*, *PABPC1*, *CPSF6* and the control LacZ gene in HEK293 cells, or of *Clp1* in the 1C6 mouse thymic epithelial cell line (Mizuochi, Kasai, Kokuho, Kakiuchi, & Hirokawa, 1992) maintained in complemented DMEM high glucose medium, was performed by infection with lentivirus-expressing shRNAs. shRNAs were cloned into the lentivirus vector pLKO with an expression driven by the ubiquitously active U6 promoter, and were provided by the RNAi Consortium of the Broad Institute, as lentiviral particles at ∼10^7^ VP/mL. For each candidate, we tested an average of 5 specific shRNAs among those with the highest knockdown efficiency as measured by the RNAi platform of the Broad Institute.

HEK293 or 1C6 cells were seeded at a density of 250,000 or 650,000 per well in 6-well plates. The next day, cells were supplemented with 8mg/ml polybrene and infected with 20 µL of shRNA-bearing lentiviruses. Each shRNA was tested in duplicate. The day after, the medium was changed to a fresh one containing 2 µg/ml puromycin. Cells were maintained in selective medium during 6 days and harvested for RNA extraction using TRIzol (ThermoFischer). Knockdown efficiency was analyzed by real-time PCR – carried out in technical triplicate – in comparing the level of expression of each candidate in the knockdown vs. control samples using the human or murine GAPDH gene for normalization with primers listed in **Supplementary File 2**. First-strand cDNA was synthesized using SuperScript II Reverse Transcriptase (ThermoFischer) and oligo(dT)12-18 primers. cDNA was used for subsequent PCR amplification using the 7300 Real-Time PCR system (Applied Biosystems) and SYBR Green Select Master Mix (ThermoFisher). Knockdown efficiency of each specific shRNA was summarized in **Supplementary File 1**. We used for subsequent analyses the extracted RNA corresponding to the 3 shRNAs yielding the highest reduction of their target mRNA (>60%).

### Lentigenic mouse generation

Three shRNAs against *Clp1* were cloned into a cluster of micro RNAs construct, the two most efficient shRNAs in two copies and the third in a single copy (**Supplementary File 1**). This construct was transferred to a lentiviral backbone, downstream of the doxycycline inducible promoter TRE3G and upstream of the ZsGreen protein. A second construct expressing the TetOn3G transactivator under the control of the EF1 promoter was generated. A single ultra-high purified and concentrated lentivector (2.2×10^9^ TU/mL) containing both constructs was generated and purified by both Tangential Flow Filtration and Chromatography. Cloning and lentiviral production were performed by Vectalys (www.vectalys.com). Fertilized oocytes (B6) were microinjected under the zona pellucida with the lentivirus suspension as described (Giraud et al., 2014). A pool of 33 transduced oocytes were reimplanted into five pseudopregnant females. Newborns were selected for integration of both constructs by PCR with primers matching the ZsGreen or TetOn3G sequences (**Supplementary File 2**). At 3 weeks of age, mice were treated with doxycycline food pellets (2 g/kg) for two weeks and then sacrificed for mTEChi isolation. Mice expressing GFP in >10% of mTEChi were selected for GFP+ and GFP-mTEChi RNA-seq.

### RNA-seq and d3’UTR ratios

Total RNA was isolated by TRIzol extraction (ThermFisher) and was used to generate polyA-selected transcriptome libraries with the TruSeq RNA Library Prep Kit v2, following the manufacturer’s protocol (Illumina). Sequencing was performed using the Illumina HiSeq 1000 machine and was paired-end (2×100bp) for mTEChi and *Aire*-KO mTEChi isolated from pooled thymi. Sequencing was single-end (50bp) for transfected or knockdown HEK293 cells, actinomycin D-treated mTEChi, and mTEChi isolated from *Clp1* lentigenic mice. RNA libraries from thymic cells isolated from lentigenic mice or actinomycin D-treated mTEChi were constructed with the Smarter Ultra Low Input RNA kit (Clontech) combined to the Nextera library preparation kit (Illumina). Paired-end (100bp) datasets were homogenized to single-end (50bp) data by read-trimming and concatenation. Lower quality reads tagged by the Illumina’s CASAVA 1.8 pipeline were filtered out and mapped to the mouse or human reference genome (mm9 or hg19) using the Bowtie aligner. For the multi-tissue comparison analysis, duplicate reads were removed with the Samtools rmdup function. For read counting, we used the intersectBed and coverageBed programs of the BEDtool distribution with the -f 1 option. It enabled the count of reads strictly contained in each exon of the Refseq genes whose annotation GTF file was downloaded from the UCSC web site (http://hgdownload.cse.ucsc.edu/goldenPath/mm9 (or hg19) /database). Differential fold-change expression between two datasets was computed using DESeq and gene expression was quantified in each sample as reads per kilobase per million mapped reads (RPKM). For d3’UTR ratio calculation, we split the last annotated exon of genes harboring a proximal pA (as identified in the PolyA_DB 2 database - http://exon.umdnj.edu/polya_db/v2/) in two distinct features and normalized the expression of the region downstream of the proximal pA to the one upstream. A proximal pA was validated when its genomic location from the PolyA_DB 2 database differs from 20bp at least to the genomic location of the UCSC annotated 3’UTR distal boundary or distal pA. In case of multiple proximal pAs, the most proximal one was considered. d3’UTR annotation files in mice and humans for RNAseq analyses are provided in **Figure 1–source data 1**.

### Multi-tissue comparison analysis

First, an RNA-seq database of mouse tissues was assembled in collecting 23 RNA-seq datasets generated from polyA-selected RNA and Illumina sequencing. The raw read sequences were obtained from the GEO database: GSE29278 for bone marrow, cerebellum, cortex, heart, intestine, kidney, liver, lung, olfactory, placenta, spleen, testes, mouse embryonic fibroblast (MEF), mouse embryonic stem cells (mESC), E14.5 brain, E14.5 heart, E14.5 limb and E14.5 liver; GSE36026 for brown adipocytes tissue (BAT) and bone marrow derived macrophage (BMDM); GSM871703 for E14.5 telencephalon; GSM879225 for ventral tegmental area VTA. Reads of the VTA dataset, over 50bp in length, were trimmed for homogeneous comparison with our RNA-seq data. We processed each collected dataset for gene expression profiling and d3’UTR ratio computing as above. To avoid center-to-center biases, we removed from the sequence assemblies the duplicate reads that could arise from PCR amplification errors during library construction. For multi-sample comparison analysis, our mTEChi datasets were also subjected to removal of duplicate reads.

Next, the tissue-specificity (one tissue of restricted expression) or selectivity (two-to-four tissues of restricted expression) of the Aire-sensitive genes was characterized by using the specificity measurement (SPM) and the contribution measurement (CTM) methods (Pan et al., 2013). For each gene, the SPM and the CTM values were dependent on its level of expression in each tissue. If no read was detected in a tissue, the latter was excluded from the comparison. For tissues of similar type, only the one showing the highest level of gene expression was included in the comparison. Cerebellum, cortex, E14.5 brain, E14.5 telencephalon and VTA referred to a group, as well as did E14.5 heart and heart, or E14.5 liver and liver. If the SPM value of a gene for a tissue is > 0.9, then the gene is considered tissue-specific for this particular tissue. Otherwise, if the SPM values of a gene for two to four tissues were > 0.3 and its CTM value for the corresponding tissues was > 0.9, then the gene was considered tissue-selective for these tissues. If these conditions were not met, the gene was left unassigned.

Finally, for the analysis of 3’UTR length variation of Aire-sensitive PTA genes between mTEChi and their tissues of expression, peripheral d3’UTR ratios of tissue-specific genes were selected in their unique identified tissues of expression. For tissue-selective genes, the d3’UTR ratios for peripheral tissues were selected randomly among their tagged tissues of expression.

### CLIP-seq analysis

The location of the 3’ end processing complex on the transcripts of Aire target genes was tracked by measuring the density of reads that map to these transcripts in PAR-CLIP sequencing data of RNAs bound to the endogenous CSTF2 from the GEO database (GSM917676). We processed these assemblies of RNA-mapped reads by the sitepro program (CEAS distribution) to infer the read density at the vicinity of proximal and distal pAs of transcripts of Aire target genes in HEK293 cells. The Aire target genes were identified from RNA-seq data of control and *Aire*-transfected HEK293 cells. The genomic location of pAs on hg19 was extracted from the PolyA_DB 2 database as above.

### Microarray gene expression profiling

Total RNA was prepared from harvested knockdown HEK293 cells using TRIzol (ThermoFisher). Single-stranded DNA in the sense orientation was synthesized from total RNA with random priming using the GeneChip WT Amplification kit (Affymetrix). The DNA was subsequently purified, fragmented, and terminally labeled using the GeneChip WT Terminal Labeling kit (Affymetrix) incorporating biotinylated ribonucleotides into the DNA. The labeled DNA was then hybridized to Human Gene ST1.0 microarrays (Affymetrix), washed, stained, and scanned. Raw probe-level data (.CEL files) were normalized by the robust multiarray average (RMA) algorithm and summarized using the R-package aroma.affymetrix (www.aroma-project.org/). d3’UTR annotation files in mice and humans for microarray analyses are provided in **Figure 1–source data 1**.

### Individual probe-level microarray analyses

For each HEK293 samples of a microarray comparison, a probe-level expression file was generated using aroma.affymetrix just before the summarization step (see above). Probes with expression values over the background in each sample, had their hg19 genomic location retrieved from data of the Affymetrix NetAffx Web site, and were mapped to the d3’UTR annotation file (generated above) using our R-implementation of PLATA (Giraud et al., 2012). d3’UTRs ratios were computed for genes having at least two individual probes in their d3’UTR regions by dividing the specific d3’UTR expression by the whole-transcript expression. We then tested between two HEK293 samples the proportion of genes with a significant increase or decrease of d3’UTR ratios to the proportion of those whose variation is not significant.

### Gene set enrichment analysis

The overlap between the transcripts impacted by Aire in HEK293 cells and those impacted by the knockdown of *CLP1*, *HNRNPL*, *DDX5*, *DDX17*, *PARP1* was tested by the GSEA software (Subramanian et al., 2005) (Broad Institute). The Aire-sensitive genes were identified from RNA-seq data of control and *Aire*-transfected HEK293 cells. For this analysis, the top 30% of genes the most sensitive to Aire were considered.

### Statistical analysis

Determination of the statistical significance differences between two experimental groups was done by the non-parametric Wilcoxon test, unless specified.

## Acknowledgments

We thank Dr Sheena Pinto for her useful comments and suggestions, that have helped improve this paper. We thank Drs. D. Mathis and C. Benoist (Harvard Medical School) for *Aire*-KO (B6) mice. We thank members of the “Homologous recombination” core facility (Cochin Institute) for lentigenic generation. This work was supported by the Agence Nationale de la Recherche (ANR) 2011-CHEX-001-R12004KK (to M.G.), the European Commission CIG grant PCIG9-GA-2011-294212 (to M.G.) and by the “Investissements d’Avenir” program managed by the ANR to the France Génomique national infrastructure ANR-10-INBS-09 (F.C.) and the French Institute of Bioinformatics ANR-11-INBS-0013 (C.B.). C.G. and Y.-C.L. were supported by fellowships from the Fondation pour la Recherche Médicale FDT20150532551 and ING20121226316, respectively. C.G., N.J. and M.G. designed the study and wrote the manuscript; C.G. and N.J. performed most of the experimental work; F.P. performed bioinformatics analyses and experiments, notably co-immunoprecipitations; F.P., Y.-C.L. and M.G. performed bioinformatics analyses; C.B. provided us with computing resources on the IFB national service infrastructure in bioinformatics and help with script optimization, O.U. and N.F. with RNA-seq and microarray datasets, and D.E.R. with lentivirus materials and editing of the manuscript.

---

**Figure 1–figure supplement 1.**
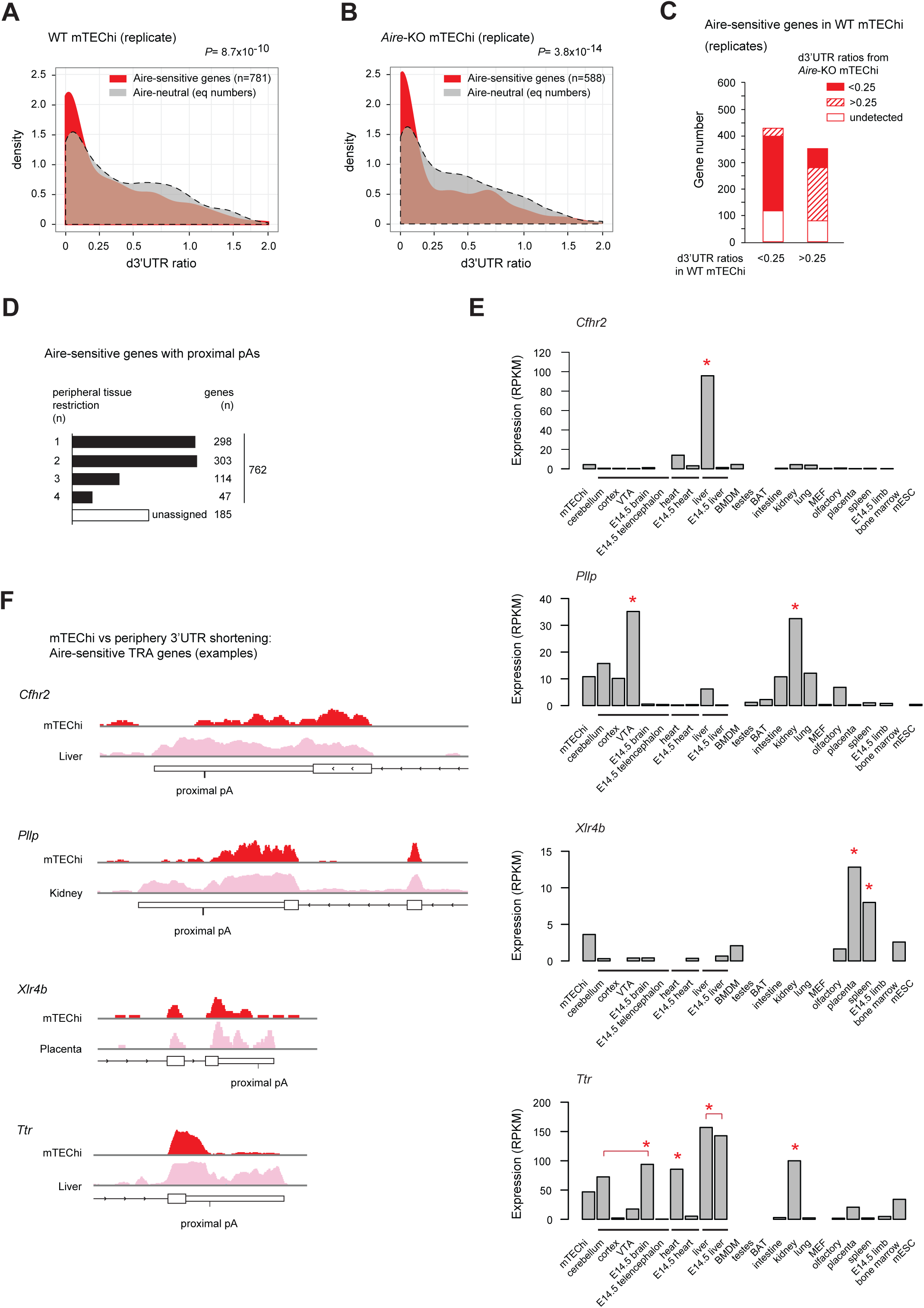
Validation and examples of the preferred short-3’UTR isoform expression of Aire-sensitive genes in mTEChi. (**A**) Densities of d3’UTR ratios of Aire-sensitive genes upregulated by Aire in mTEChi and of Aire-neutral genes from a replicate RNA-seq experiment in WT and *Aire*-KO mTEChi sorted from a pool of 4 thymi; equal number (n=781) of selected neutral genes, asinh scale. (**B**) Densities of d3’UTR ratios of Aire-sensitive and neutral genes in *Aire*-KO mTEChi; replicate experiment, equal number (n=588) of selected neutral genes, asinh scale. (**C**) Proportion of Aire-sensitive genes with d3’UTR ratios <0.25 or >0.25 in *Aire*-KO mTEChi among those with d3’UTR ratios <0.25 or >0.25 in WT mTEChi; replicate experiment. (**D**) Number of Aire-sensitive TRA genes, *i.e.*, specific or selective of two to four tissue types across 16 groups of tissues of similar type from 22 collected mouse-tissue RNA-seq datasets. (**E**) Level of expression of *Cfhr2*, *Pllp*, *Xlr4b* and *Ttr* (taken as examples of Aire-sensitive TRA genes) from RNA-seq data of mTEChi and 22 mouse tissues. BAT stands for brown adipocytes tissue, BMDM for bone marrow derived macrophage, MEF for mouse embryonic fibroblast, mESC for mouse embryonic stem cells and VTA for ventral tegmental area. Tissues of similar types are binned together (black line). *Cfhr2* is specific to the liver; *Pllp* is selective to the brain and kidney; *Xlr4b* to the placenta and spleen and *Ttr* to the brain, heart, liver and kidney. (**F**) Examples of Aire-sensitive TRA genes with 3’UTR shortening in mTEChi. Annotated 3’UTRs are represented by thin boxes. For each gene, the mapped RNA-seq reads are shown in mTEChi (red) and in its tissue of normal expression (pink).

**Figure 2–figure supplement 1.**
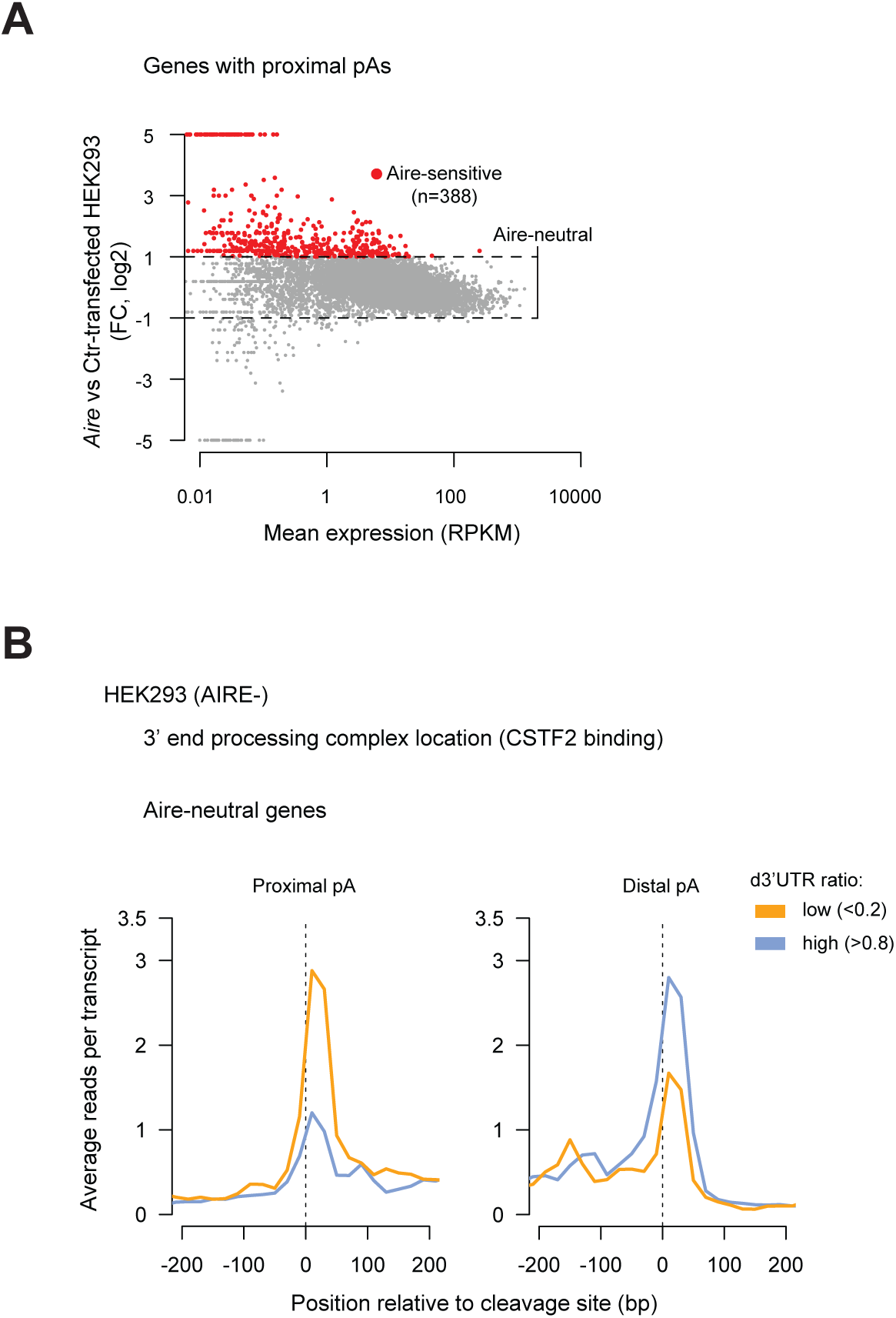
Correlation of the binding of the 3’ end processing complex with proximal pA location. (**A**) RNA-seq differential expression (fold-change) of genes with proximal pAs between *Aire*-transfected and Ctr-transfected HEK293 cells. Red dots show genes upregulated by twofold or more (Z-score criterion of *P*<0.01) (Aire-sensitive). Genes between the dashed lines have a change in expression less than twofold (Aire-neutral). (**B**) Average density of reads from PAR-CLIP analyses in HEK293 cells (AIRE-) of CSTF2 protein as a marker of the 3’ end processing complex, in the vicinity of proximal and distal pAs of Aire-neutral genes with high d3’UTR ratios > 0.8 or low d3’UTR ratios < 0.2.

**Figure 3–figure supplement 1.**
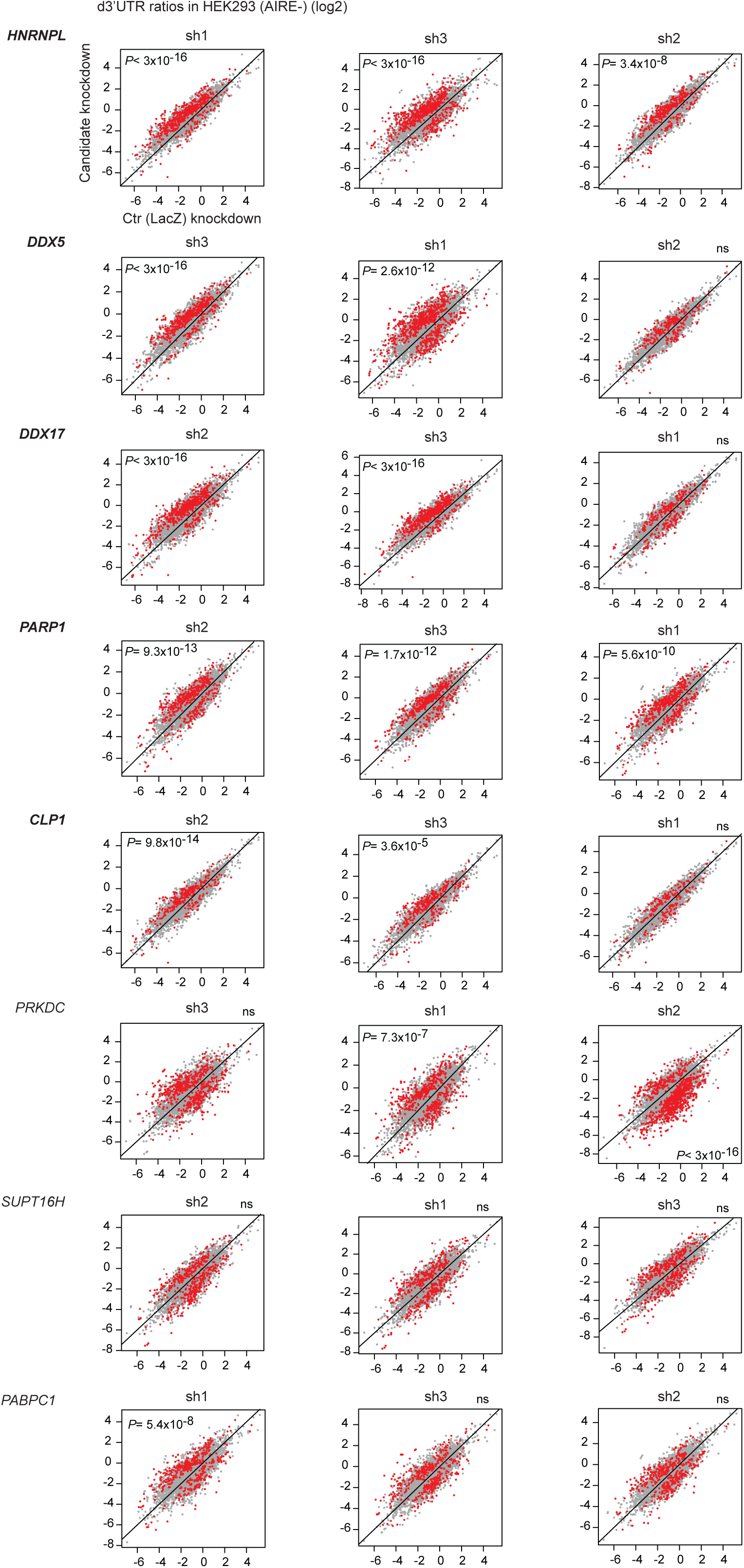
Effect of shRNA-mediated interference of candidate factors on d3’UTR ratios. Individual probe-level analysis of microarray data from HEK293 cells infected by lentiviruses containing one of the three hit shRNAs of each candidate gene or the Ctr (LacZ) shRNA. Genes whose d3’UTR ratios vary significantly from a candidate KD sample to the Ctr sample are shown in red, otherwise in gray. P values comparing the proportion of genes with a significant increase or decrease of d3’UTR ratios to the proportion of genes whose variation is not significant, are assessed by a Chi-squared test and labeled in the quadrant toward which the d3’UTR ratios (in red) significantly increase.

**Figure 3–figure supplement 2.**
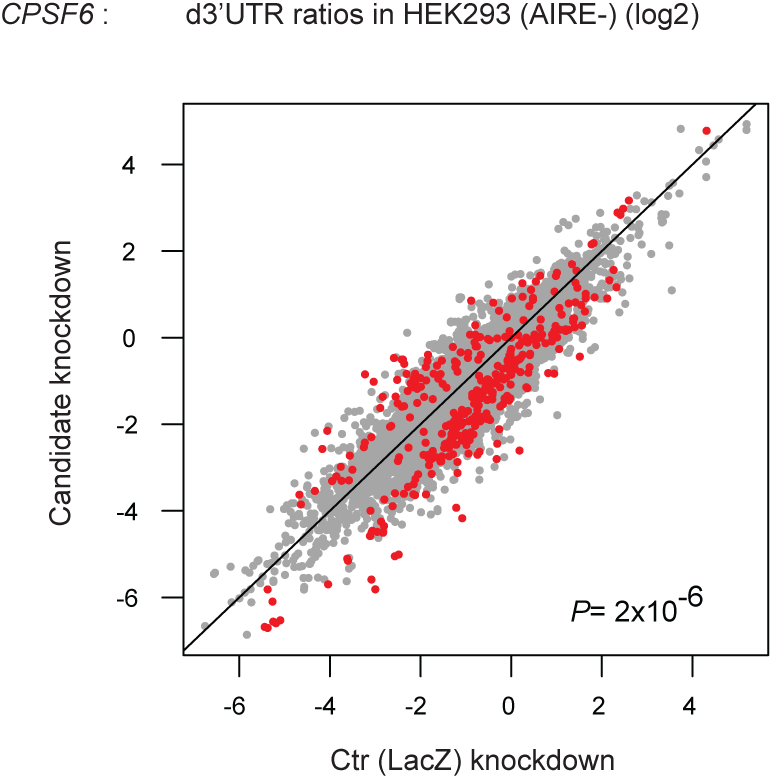
CPSF6 promotes 3’UTR lengthening. Individual probe-level analysis of microarray data in HEK293 cells infected by a lentivirus targeting *CPSF6*. Genes whose d3’UTR ratios vary significantly from the *CPSF6* KD sample to the Ctr (LacZ) KD sample are shown in red, otherwise in gray.

**Figure 4–figure supplement 1.**
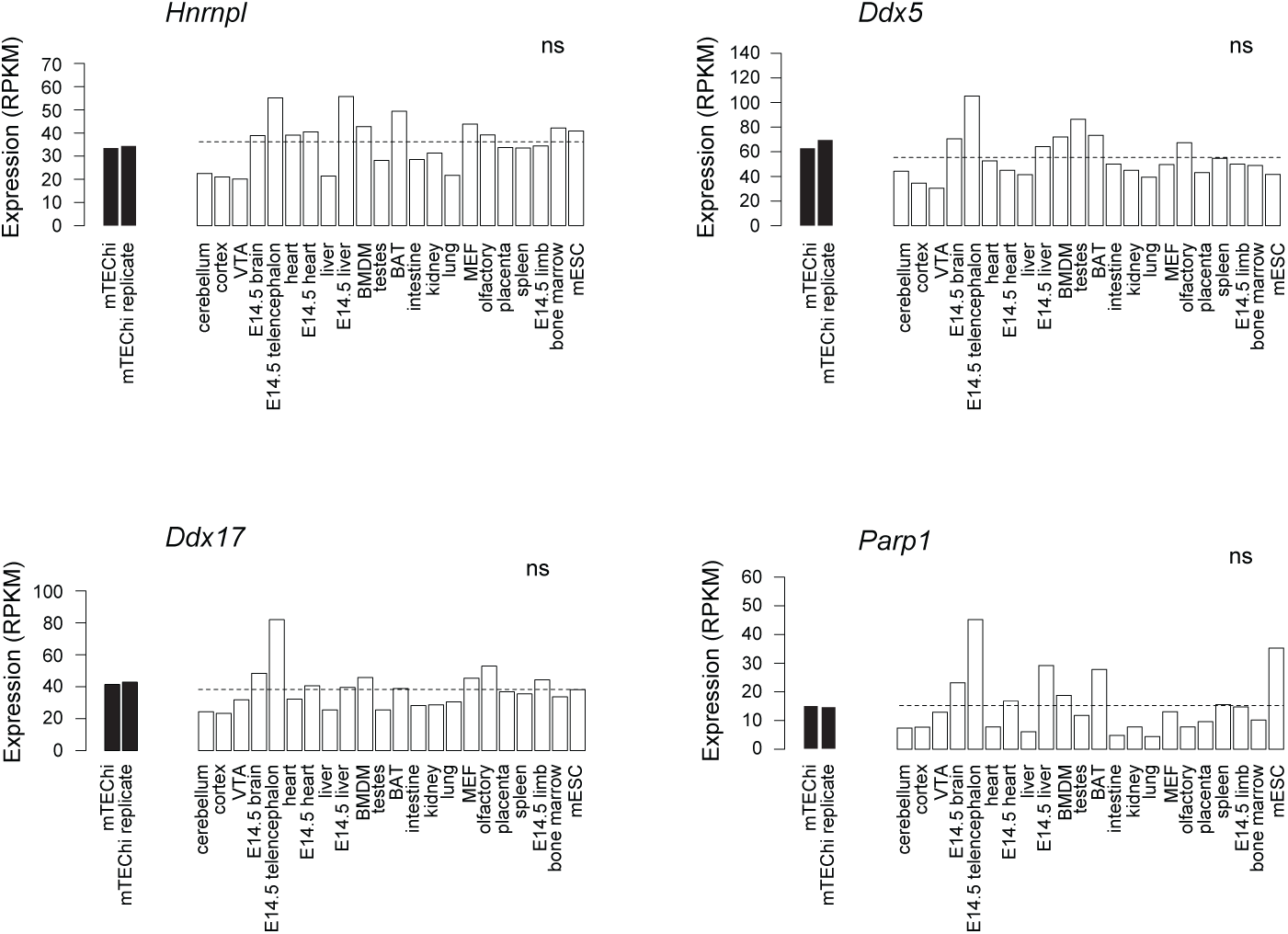
Comparison of gene expression in mTEChi and in mouse tissues. Expression of the candidate factors having an effect on 3’UTR shortening in HEK293 cells from RNA-seq data of two replicate mTEChi samples, and 22 mouse tissues. The dashed lines show the median expression of the factors in the tissues.

**Figure 4–figure supplement 2.**
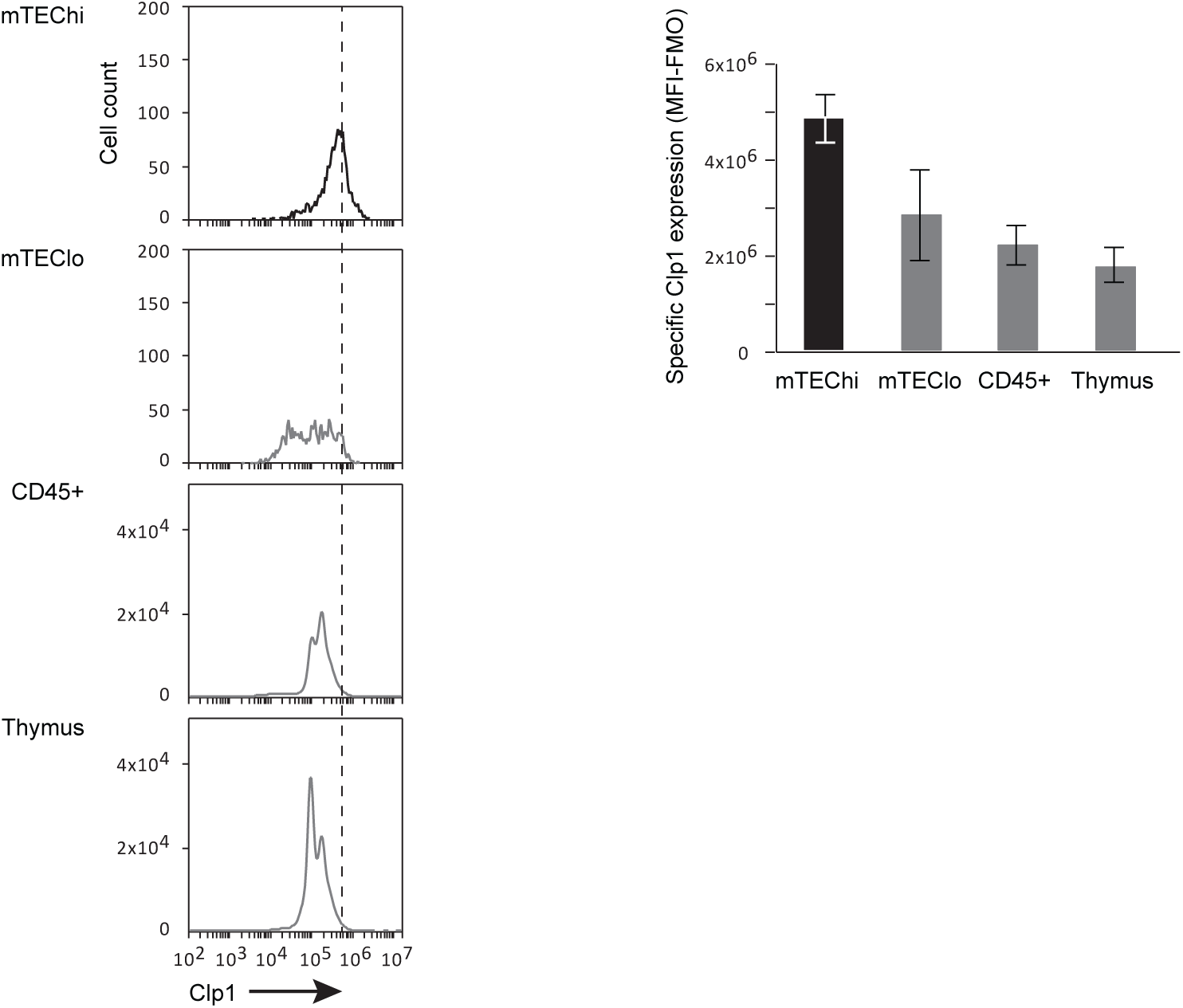
Clp1 expression in the thymus. Clp1 expression is shown as Mean Fluoresence Intensity (MFI) in mTEChi, mTEClo, thymic-sorted CD45+ and the entire thymus (*Left*) and is subtracted of fluorescence minus one (FMO) control signals from two independent experiments (*Right*).

**Figure 4–figure supplement 3.**
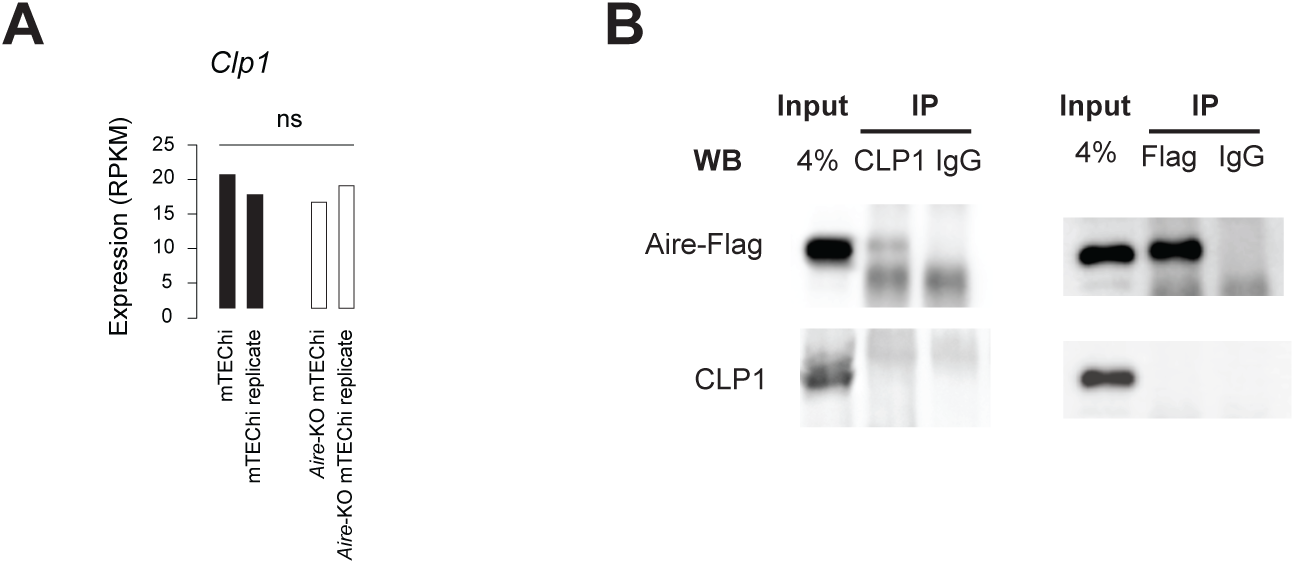
Clp1 is not linked to nor controlled by Aire. (**A**) Clp1 expression levels are neutral to *Aire* deletion. (**B**) Coimmunoprecipitation of endogenous CLP1 with Flag-tagged Aire is not specific to CLP1, the latter been not detected in the anti-CLP1 immunoprecipitate. No interaction is detected in the reciprocal immunoprecipitation.

